# One Health genomic epidemiology reveals cryptic transmission and surveillance gaps in a decade-long urban rabies epidemic in Arequipa, Peru

**DOI:** 10.64898/2026.07.07.736726

**Authors:** Kirstyn Brunker, Alejandra Dávila-Barclay, Elvis W. Díaz, Sandeep Kasaragod, Rowan Durrant, Katie Hampson, Edith Zegarra, Ynés Monroy, Ricardo Castillo-Neyra

## Abstract

Persistent local foci remain a major barrier to eliminating dog-mediated rabies, yet the processes sustaining transmission at micro-scales, particularly within complex urban systems, remain poorly understood. We apply an integrated One Health genomic epidemiology framework to reconstruct a decade-long rabies virus (RABV) epidemic in Arequipa, Peru. Combining 133 newly generated whole genomes with existing data, we assembled the most comprehensive canine RABV dataset in Latin America and used whole-genome-informed phylogenetic, phylodynamic and landscape analyses to trace the epidemic from its first detection in 2015. Transmission was dominated by a single lineage, estimated to have been introduced around 2012, which circulated for approximately three years before detection. We estimate that only 1–2% of infections are routinely detected, revealing extensive cryptic transmission and highlighting the limitations of case-based metrics for verifying disease freedom. Although additional regional and transboundary introductions were identified, only one resulted in sustained transmission. Within Arequipa city, transmission was highly structured, concentrated in densely populated and socioeconomically deprived areas, and shaped by urban connectivity. Roads and dry water channels facilitated spread, while rivers acted as partial barriers. Our findings demonstrate that rabies persistence reflects interacting processes across spatial scales, from regional introductions to highly structured urban transmission. They highlight the importance of genomic-informed, spatially targeted surveillance and control strategies to identify and address persistent transmission foci and accelerate progress towards elimination.

**Funding:** US National Institute of Allergy and Infectious Diseases, US Fogarty International Center, Wellcome Trust, UK Medical Research Council, and UK Engineering and Physical Sciences Research Council.

## Introduction

Rabies is a fatal zoonosis responsible for tens of thousands of human deaths annually, despite effective vaccines for both humans and animals^1^. Most human cases result from dog-mediated transmission, and global efforts aim to end human deaths from canine rabies by 2030^2^. Mass dog vaccination has drastically reduced disease burden across Latin America but elimination remains incomplete, with persistent transmission in several regions and continued risk of re-emergence in areas declared rabies-free^3^.

Urban and peri-urban rabies virus (RABV) persistence is a particular challenge. Unlike at broader spatial scales, where viral populations are structured by historical colonisation events, geographic barriers and host associations^4,5^, urban environments combine high host density, heterogeneous infrastructure, and frequent human-mediated movement, producing co-circulating lineages and overlapping transmission chains^6,7^. These dynamics sustain cryptic endemic circulation, and conventional surveillance– which relies on the timing and location of detected cases– gives only a partial view of the mechanisms behind persistence.

Whole-genome sequencing (WGS) and genomic epidemiology have transformed the study of RABV, showing how distance and landscape barriers structure spread at regional and continental scales^4,5,8^ and how contact networks, population density, and reintroductions shape it at finer scales^6,7,9–11^.

However, few studies have resolved transmission at the spatio-temporal scales needed to inform urban rabies control, and translation into actionable surveillance remains limited.

This gap matters most where elimination is within reach, as in Latin America. As incidence falls, surveillance must detect low-level persistence, identify sources of re-emergence and distinguish local transmission from repeated introductions, questions that are hard to answer with epidemiological data alone. Sequence data, integrated with spatial, environmental, and intervention data within a One Health framework^12,11,9^, and as an independent line of evidence, can reconstruct transmission histories^5,13^, identify introductions^10,12,11,9^, verify elimination^11^, and estimate epidemic size^14^. Yet, genomic surveillance is largely absent from routine rabies control across Latin America, with only one study (our previous work) publishing contemporary WGS data^10^.

Arequipa, Peru, exemplifies this gap. The city and region were included in the 2013 REDIPRA declaration that 88% of Peruvian territory was free of dog-mediated rabies, following 14 years of epidemiological silence^15^. Yet, transmission re-emerged in the region in 2014 and in the city in early 2015^16^ (Fig. 1). Over the subsequent decade, approximately 400 laboratory-confirmed canine rabies cases were detected despite mass dog vaccination and reactive outbreak response, albeit at varying frequency^17^. Epidemiological and One Health investigations identified spread along urban water channels that are dry for most of the year^18^, expansion into peri-urban areas, and associations with socio-economic status^19^. However, these inferences derive from surveillance and modelling, which cannot alone disentangle the processes generating epidemic patterns. Key uncertainties therefore remain, including the timing and source of introduction(s), the extent of undetected transmission, the relative importance of local versus long-distance spread, and how control can be optimised to interrupt transmission.

**Fig. 1.**
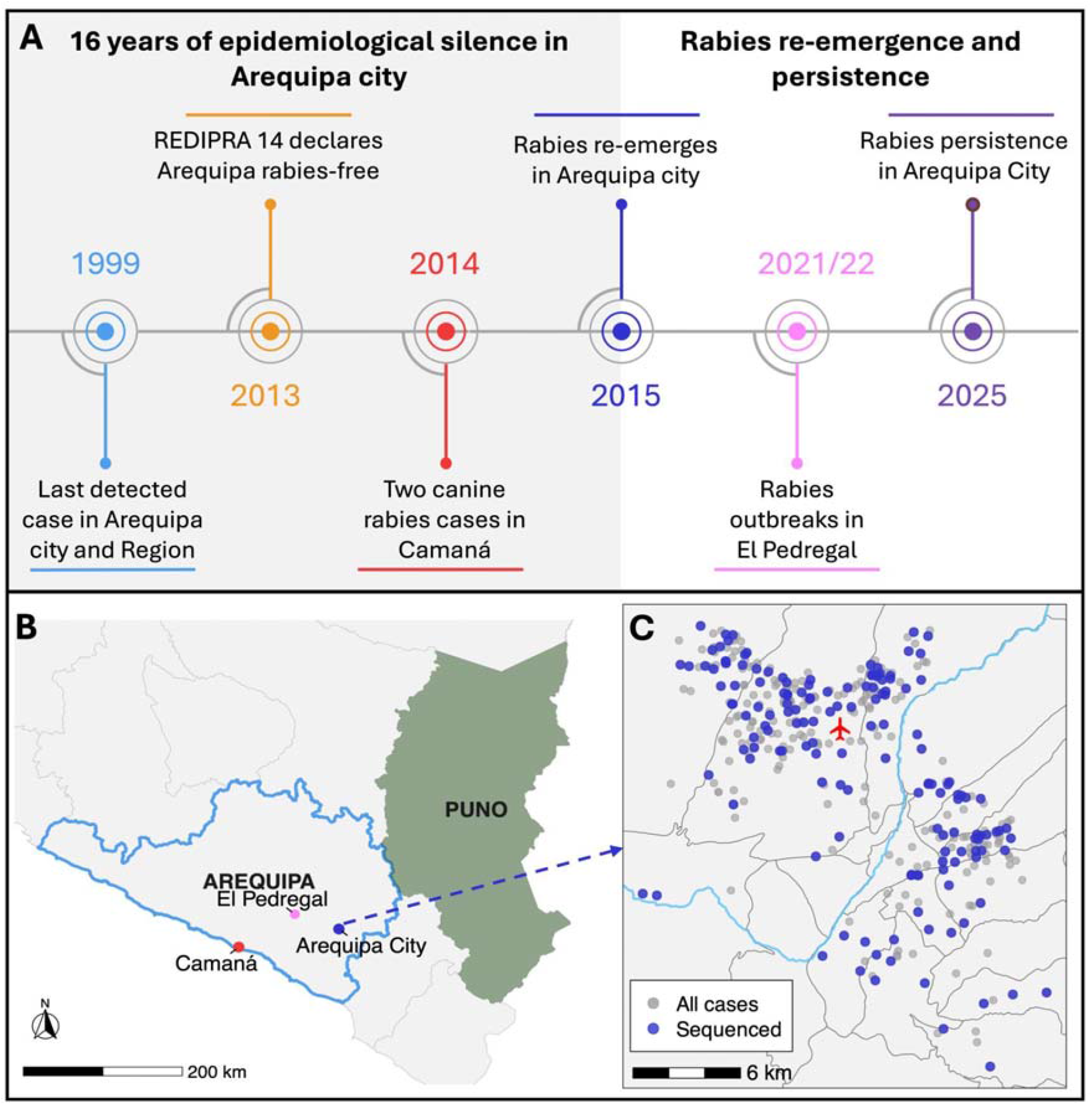
Epidemiological timeline and spatial context of rabies elimination, reintroduction, and persistence in Arequipa, Peru. (A) Timeline of canine rabies in Arequipa showing 16 years of epidemiological silence following the last detected case in 1999 and declaration of rabies-free status in 2013, followed by re-emergence in 2014–2015 and persistence through to 2025. (B) Map of the study area showing Arequipa region (blue outline) and key locations implicated in rabies events. (C) Spatial distribution of reported rabies cases in Arequipa city, with the Chili River and Rodríguez Ballón International Airport indicated for spatial context. Grey points show all reported cases; blue points indicate sequenced samples, with grey polygons indicating administrative districts.

In this study, we applied an integrated genomic epidemiology framework to reconstruct a decade-long canine rabies epidemic in Arequipa, Peru, providing insights into RABV origins, transmission dynamics, and the mechanisms underlying viral persistence.

## Methods

The study protocol (No. 216867) was approved by the Institutional Ethics Committee for the Use of Animals (IECA) of the Universidad Peruana Cayetano Heredia (CERTIFICATE-IECA-021-04-25).

### Study area and surveillance

The city of Arequipa is located in the Southern Peruvian Andes within the Arequipa Region, bordering the Puno Region, which shares a boundary with Bolivia. Arequipa city comprises a central urban core surrounded by expanding peri-urban settlements arranged in a ring of younger neighbourhoods. The Chili River bisects the city into north-west and south-east sectors, linked by multiple pedestrian and vehicle bridges, with an associated water channel system extending throughout the urban area. Following 14 years without cases, the region was declared rabies-free in 2013 but canine rabies re-emerged in Camaná in 2014 and reached Arequipa city in March 2015 (Fig. 1), spreading to nearly all city districts by March 2025 with occasional transmission to neighbouring El Pedregal. Control relied primarily on mass dog vaccination with variable coverage, alongside outbreak response (ring vaccination, case investigation, contact tracing).

Between 2015 and 2025, 398 animal cases were reported (386 in Arequipa city: 383 dogs, 3 cats; 12 dogs in El Pedregal), and one human death in 2023. Surveillance is overseen by the Arequipa-Caylloma Health Network, which submits brain samples to the Arequipa Regional Reference Laboratory (LRR). The LRR also receives samples from private clinics, veterinary professionals, and the research team. Cases were confirmed by direct fluorescent antibody (DFA) testing, with duplicates confirmed at the Peruvian National Institute of Health and discrepancies resolved by LN34 RT-PCR^20^ or mouse inoculation where available.

### Sample processing and sequencing

DFA-positive brain samples in DNA/RNA Shield were provided by the LRR, with epidemiological metadata supplemented by outbreak-response case investigations (dates, district, travel history and coordinates). Total RNA was extracted using the Quick-RNA Viral kit (Zymo Research, Irvine, CA, USA), reverse-transcribed and amplified by RT-PCR with a Peru-specific RABV primer scheme^10^, following established sample-to-sequence protocols^21–24^. Libraries (Oxford Nanopore SQK-NBD114.24, with negative controls in each run) were sequenced on R10 flow cells on a MinION Mk1B, and consensus sequences generated using an established pipeline^22^. In total, 133 genomes were generated and combined with previously published Peruvian WGS, giving 167.

### Phylogenetic and phylodynamic analyses

Analyses were conducted across three nested spatial scales.

#### Regional

Publicly available Latin America and Caribbean (LAC) canine RABV sequences (>200 bp) and a Cosmopolitan Africa 4 outgroup (GenBank: KF154998) were retrieved from the RABV genome database (RABV-gDB; https://rabv-gdb.cvr.gla.ac.uk/) and aligned with the newly sequenced genomes (n = 133) in MAFFT v7.520. A maximum-likelihood phylogeny was inferred in IQ-TREE (GTR+Γ, 1000 ultrafast bootstraps). In R version 4.5.2, outliers (top 1% longest terminal branches) were removed using the ape package, and the tree was outgroup rooted and visualised (ggtree, sf, rnaturalearth).

#### Country

A time-scaled Bayesian phylogeny was inferred in BEAST v10.5.0^25^ using WGS from Arequipa city (n = 152), El Pedregal (n = 11), and Puno and Madre de Dios (n = 4). Initial exploratory runs identified a divergent sequence associated with an external introduction (219_2024), which was excluded from the final analysis due to its disproportionate influence on temporal structure (final n = 166). Temporal signal was confirmed by root-to-tip regression in Clockor2^26^ (Fig. S1). We used a GTR+Γ model partitioned into coding and non-coding regions, an uncorrelated lognormal relaxed clock (UCLD), and a constant population size prior. MCMC chains ran for 30 million iterations (sampled every 3,000), and trees were summarised using HIPSTR with 10% burn-in. Convergence was assessed in Tracer for all BEAST analyses (ESS > 200).

#### City

Analyses focused on the dominant local lineage (n = 148). Discrete trait phylogeography used an asymmetric discrete trait model with Bayesian stochastic search variable selection (BSSVS) under a UCLD clock, with Markov jumps (MJ) and rewards (MR) quantifying transition rates and within-state persistence. Traits were position relative to the Chili River, urbanicity (defined from locality characteristics and expert knowledge), and administrative district (INEI shapefiles; https://ide.inei.gob.pe/#capas). Because case assignment follows the local microred (health network), some cases fall geographically within one district but are assigned to another; recorded district labels were retained. Robustness to sampling bias was assessed by downsampling to n = 100 using Treemmer, reducing districts with >10 sequences (Cerro Colorado, Mariano Melgar, Yura).

Continuous phylogeography used case coordinates in BEAST (GTR+Γ, UCLD clock, Skygrid prior; 200 million iterations sampled every 20,000). Branch-specific dispersal vectors from 100 post-burn-in trees were extracted in SERAPHIM 2.0^27^ to estimate mean and weighted branch velocity and diffusion coefficients.

Landscape analyses used the same dispersal vectors on a 200 m grid covering the sequenced cases plus a 1 km buffer. Layers (rivers, roads, water channels, built-up areas, socioeconomic status (SES); Fig. S2) were converted to resistance surfaces (normalised 0–1, scaled k = 10, 100 or 1000) and tested as both resistance and conductance against a null surface of homogeneous space (details in Supplementary text). SES was represented in two ways: a continuous gradient derived from categorical classes (A–F) and rescaled (6–1) to reflect decreasing socioeconomic status, and a binary classification distinguishing affluent versus deprived areas. Linear features (rivers, roads, and water channels) were represented as binary surfaces, where influence is restricted to the mapped feature, and as distance-decay surfaces, where influence decreases with distance in 100 m intervals, capturing spatially graded infrastructure effects. SERAPHIM’s Q statistic tested whether each variable improved on a null model of geographic distance alone (significance via randomisation of lineage locations), and the E and R statistics used to assess, respectively, whether lineages are associated with particular environmental conditions and whether they move toward or away from environmental gradients. Support was summarised as approximated Bayes factors.

## Outbreak size estimation

Outbreak size and annual cases for Arequipa city were estimated following Durrant et al. (2026) by combining the total tree length of the city-scale tree (n = 148) with per-generation substitution rates from BEAST posteriors and a mean generation interval of 32 days. The proportion of cases sequenced (*p*) was obtained by solving the quadratic:

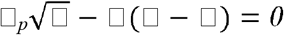

where *T* is the tree length, μ the per-generation substitution rate, and *S* the number of tips. Outbreak size was estimated as *S/p*, across 9,002 posterior rate samples. Annual incidence was derived by pruning the tree sequentially by year and differencing cumulative estimates.

## Results

### Genomic context and regional connectivity

We generated 133 new RABV WGS from Arequipa city, of which 129 achieved >90% coverage and four achieved >80% (Table S2). Combined with 34 published Peruvian genomes, this gave 167 WGS: 152 from Arequipa city, 11 from El Pedregal, 3 from Puno, and 1 from Madre de Dios. Including our sequences, the LAC dataset comprised 1,816 RABV sequences (>200 bp; Fig. S3), of which only 11% were WGS (n = 199, of which 84% are the Peruvian genomes); the rest were partial or single/multi-gene sequences.

All sequences from this study fell within the Cosmopolitan AM5 lineage. AM5 accounted for 296 of the 1,816 LAC sequences (including those from this study). Within AM5, Arequipa sequences were most closely related to those from Bolivia and Puno, indicating connectivity across the Andean corridor (Fig. 2). Bolivian sequences frequently occupied basal positions, consistent with unsampled diversity and a possible regional source population. Beyond the dominant Arequipa cluster, we identified multiple independent introductions, including a recent case clustering with sequences from Argentina, Bolivia, and Brazil (highlighted in Fig. 3), and another associated with Puno (resolved in Fig. 3; collapsed in Fig. 2).

**Fig. 2.**
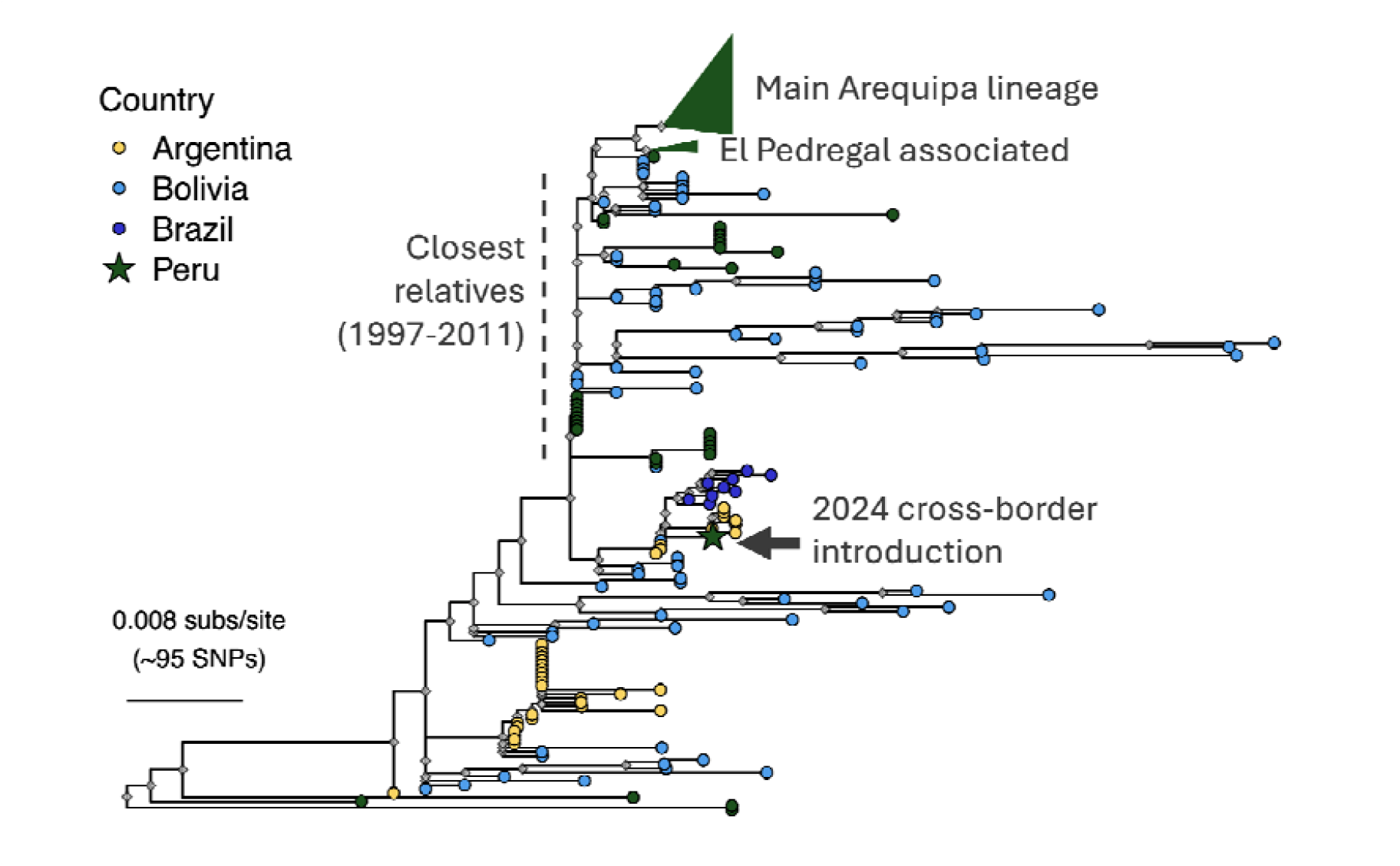
Cosmopolitan AM5 canine RABV across Latin America and the Caribbean (LAC). Maximum-likelihood phylogeny of 296 AM5 sequences from LAC, including 167 from this study, showing the placement of Arequipa viruses within a regional diversity. The dominant Arequipa and El Pedregal-associated lineages are collapsed (triangles), and a cross-border introduction to Arequipa is marked with a star. Tip colours denote country of origin. Grey diamonds indicate nodes with ≥90% bootstrap support.

**Fig. 3.**
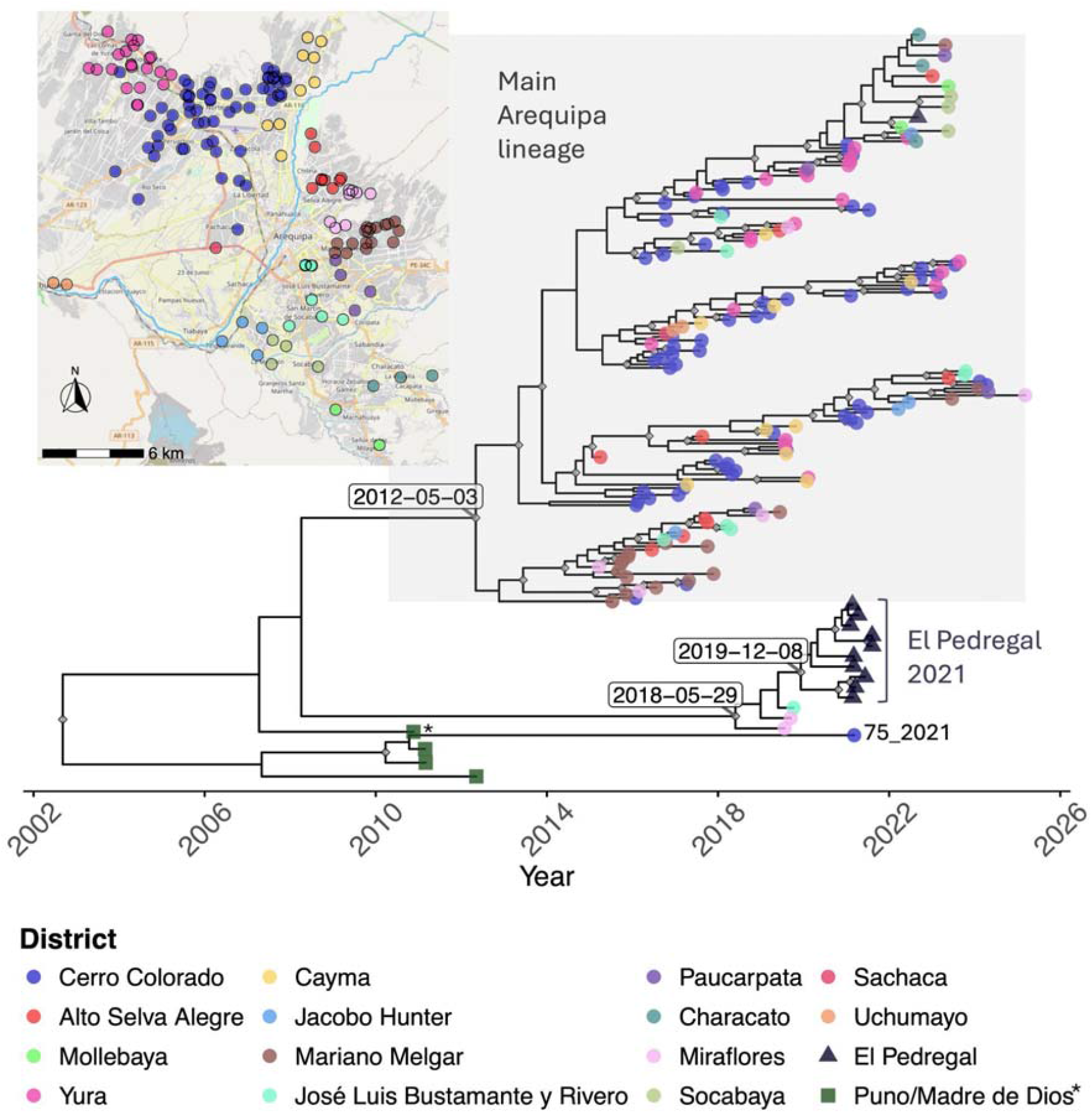
Time-scaled phylogeny of RABV genomes from Arequipa and elsewhere in southern Peru. HIPSTR (highest independent posterior subtree reconstruction) tree of genomes from Arequipa city (n = 152), El Pedregal (11), and historical isolates from the Puno (3) and Madre de Dios* (1) regions. Branch lengths are scaled in years; internal node symbols indicate posterior support > 0.9. Tip colours denote sampling location, the dominant lineage is highlighted, and sample 75_2021 is labelled as an independent introduction linked to Puno. The inset map shows sample locations within Arequipa coloured by district; apparent mismatches between location and district reflect assignment by microred (health network).

### A single dominant lineage drives sustained transmission amid repeated introductions

Time-scaled phylogenetic reconstruction showed that transmission in Arequipa is overwhelmingly dominated by a single lineage, comprising 147 of 152 Arequipa city genomes (97%), plus one 2022 case from El Pedregal (Fig. 3). This lineage was inferred to have emerged around April 2012 (95% HPD: 2011.39–2013.85), approximately three years before the first detected case in 2015, and showed limited within-cluster diversity (mean patristic distance: 13 SNPs), consistent with sustained local transmission following a single introduction. The remaining Arequipa-associated sequences comprised two independent introductions that did not persist long term, and three sequences associated with the contained 2021 El Pedregal outbreak (Fig. 3).

A distinct, well-supported lineage (n = 10) was associated with the 2021 El Pedregal outbreak, with an MRCA dated to August 2019 (95% HPD: 2019.74–2020.71). Three Arequipa sequences from 2019 were basal to this clade and, when included, moved the MRCA to May 2018. This is consistent with introduction from unsampled regional diversity, with possible transient circulation in Arequipa city before spreading to El Pedregal. The single 2022 El Pedregal case clustered within the dominant Arequipa lineage, supporting the previous conclusion of an independent introduction rather than continuation of the 2021 outbreak^10^.

Two sequences (75_2021 and 219_2024) were highly divergent from both the dominant lineage and El Pedregal lineages, supporting independent introductions. Sequence 219_2024 differed by 161– 182 SNPs from the dominant and El Pedregal lineages, while sequence 75_2021 differed by 18–39 SNPs, well above within-lineage diversity (Arequipa: 13 SNPs; El Pedregal: 8 SNPs).

### Population dynamics of the dominant lineage

Bayesian Skygrid analysis of the dominant Arequipa lineage (n = 148) showed that effective population size (N1.τ) increased between 2015 and 2017 and remained elevated thereafter, consistent with established endemic transmission (Fig. 4b). A peak around 2020 was followed by a temporary decline coinciding with the COVID-19 pandemic, and a resurgence around 2023. N1.τ then appeared to plateau, broadly consistent with reductions in reported cases. Credible intervals widen towards the present, so recent estimates are uncertain, but N1.τ remained above early-epidemic levels with no clear evidence of a sustained decline.

**Fig. 4.**
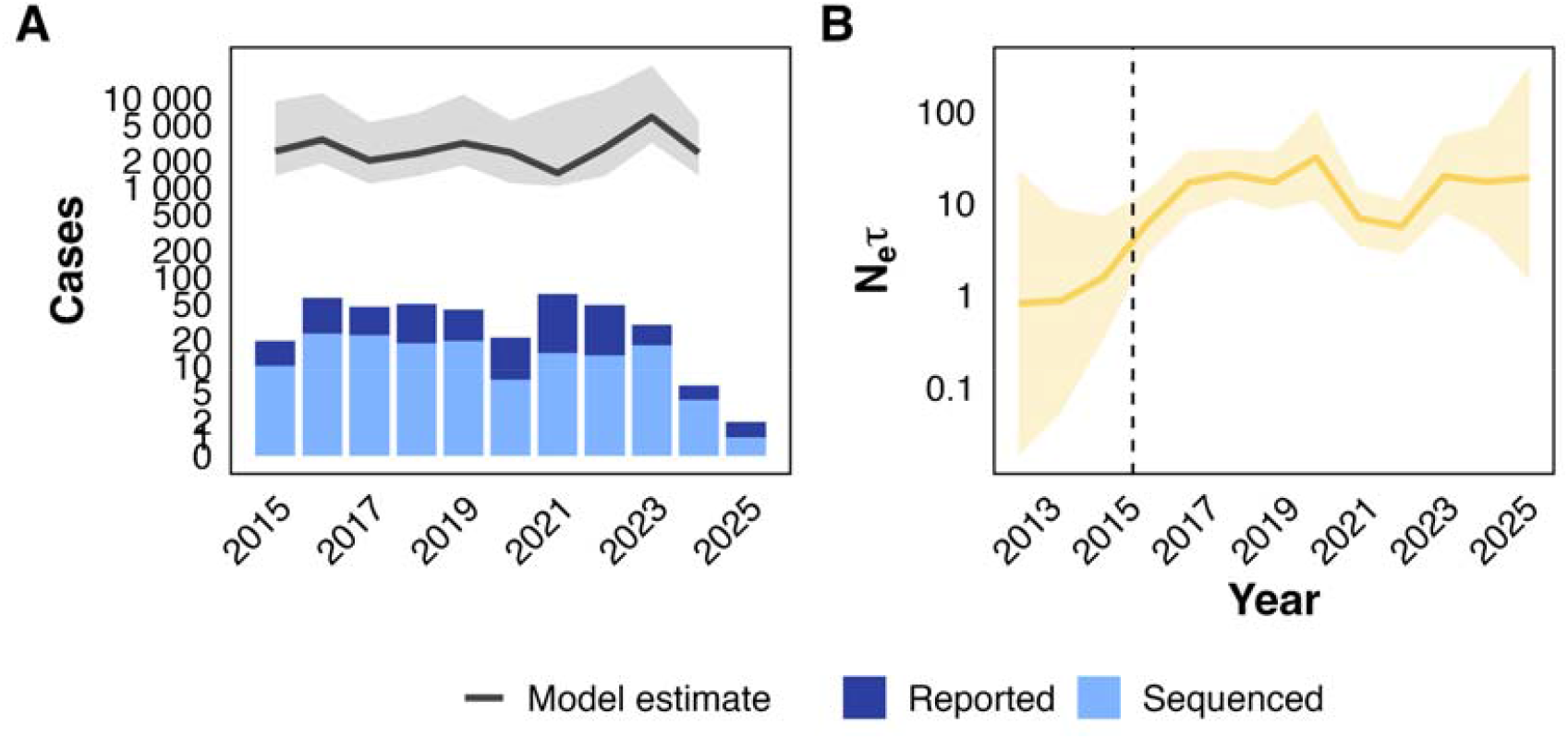
Phylogenetically estimated outbreak size and effective population size of canine rabies in Arequipa. (A) Reported (dark blue) and sequenced (light blue) cases, with phylogenetically estimated outbreak size (grey line) on a pseudo-logarithmic scale; 2025 is excluded because only one case was sequenced. (B) Bayesian skygrid estimate of effective population size (Nlilτ; line, median). Ribbons show 95% credible intervals, and the dashed line marks the first detected case.

### The epidemic is substantially under-detected

Outbreak size estimates indicate that the true scale of the epidemic in Arequipa far exceeds surveillance data. The total outbreak size was estimated at approximately 28,637 cases, implying that routine surveillance captured only 1-2% of cases annually (Fig. 4a). Annual estimates from sequentially pruned trees were affected by long branches, likely reflecting unsampled diversity with deep ancestral origins, but estimates from the full phylogeny are robust to this because they integrate unsampled diversity across the whole tree.

### Persistent local transmission is concentrated in a few hubs with directional spread

Continuous phylogeographic reconstruction of the dominant lineage (Fig. 6A) estimated a mean weighted diffusion rate of 4.15 km/year (95% HPD: 3.63–4.68 km/year) and a weighted diffusion coefficient of 6.82 km^2^/year (95% HPD: 5.72–7.72 km^2^/year). Unweighted estimates were higher (8.09 km/year; 8.40 km^2^/year), suggesting episodic long-distance dispersal alongside local transmission. A weak isolation-by-distance signal (r ≈ 0.19–0.20) indicated modest spatial structuring.

Discrete phylogeography (Fig. 5) showed strongly asymmetric movement, with specific districts acting as net sources and others as net recipients. A few preferential routes (e.g., Cerro Colorado to Yura and to Cayma) accounted for a disproportionate share of movement, while reverse transitions were rare (e.g., Yura to Cerro Colorado: 3-4 MJ). Cerro Colorado was the dominant hub (Figs. 6A, 7), accounting for 51% of total evolutionary time and the most outgoing movements (38 MJ). Of 17 transitions with BF > 10, only a subset contributed substantially to spread, indicating that transmission is concentrated along frequently used routes. Most were robust to downsampling (Table S3), although 3 (all BF < 20) were sensitive. All supported transitions are in Table S4.

**Fig. 5.**
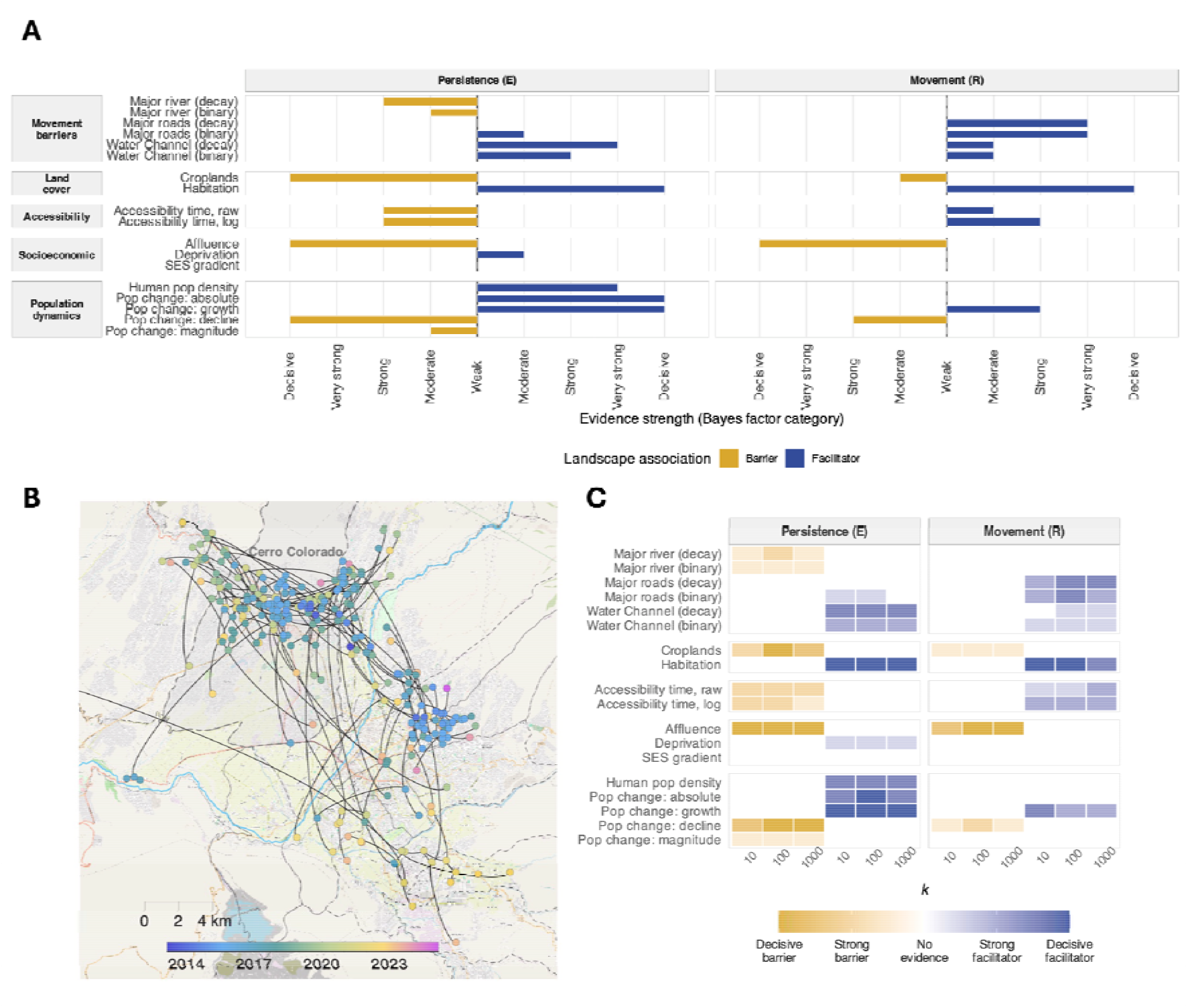
Association of landscape variables with rabies virus diffusion in Arequipa city. (A) Bars show the strength of evidence (approximated Bayes factor category) for each variable’s association with lineage persistence (E) and movement (R) in SERAPHIM analyses of the dominant lineage (n = 148), at resistance scaling factor k = 100. Bars to the left of zero (gold) indicate barriers and bars to the right (blue) facilitators. (B) Continuous phylogeographic reconstruction (relaxed random walk) of the dominant Arequipa lineage (n = 148), shown as the HIPSTR tree on the map. District boundaries are dashed and Cerro Colorado is shaded grey. Basemap: © OpenStreetMap. (C) Tiles show support for barrier (yellow) and facilitator (blue) associations at k = 10, 100, and 1000, for persistence (E) and movement (R) models. Colour intensity indicates evidence strength; white denotes weak or no evidence (BF <3).

**Fig. 6.**
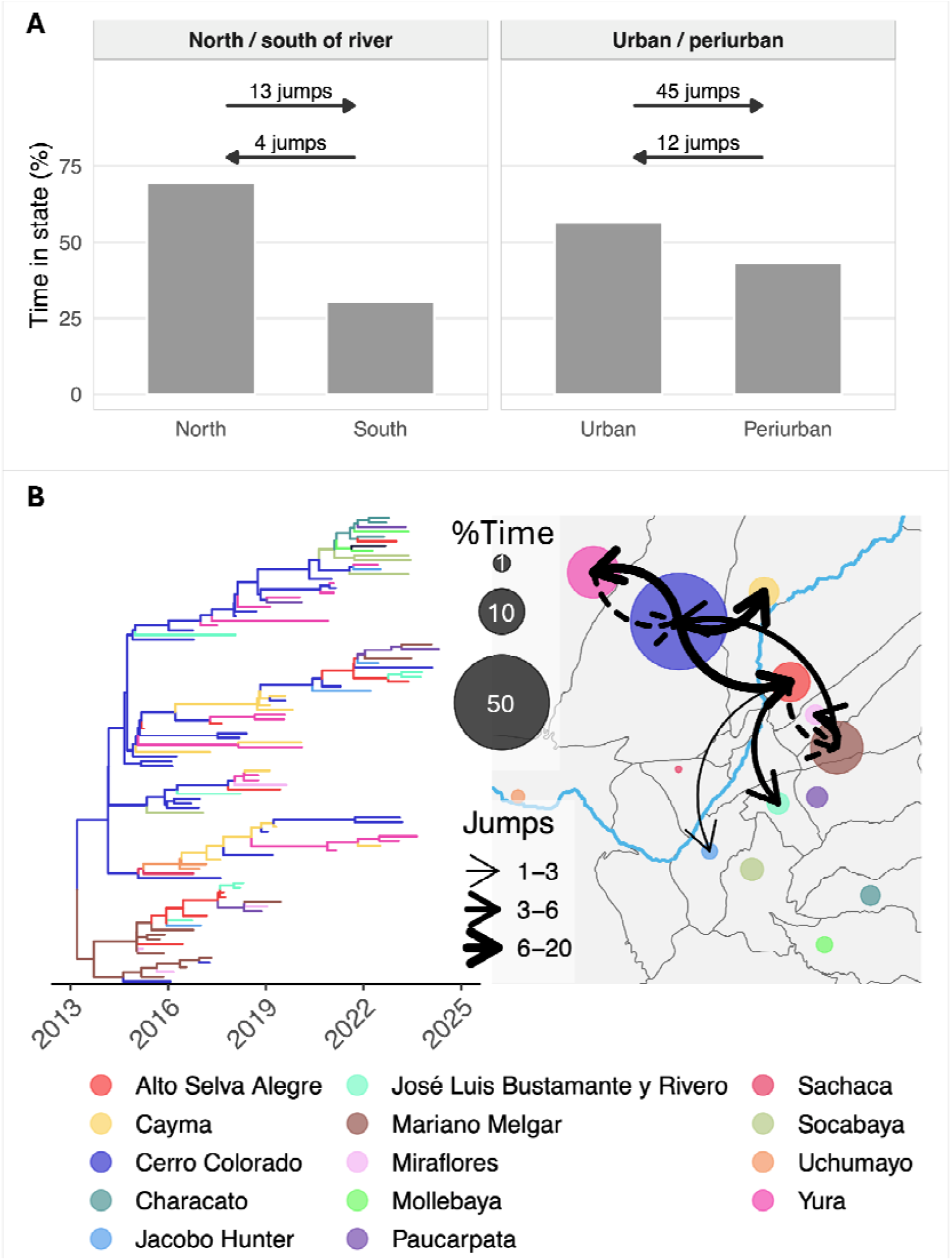
Discrete trait phylogeographic reconstruction of rabies virus transmission in Arequipa. A) Time spent in each state (bars; Markov reward, % of evolutionary time) and mean Markov jumps between states (arrows; values rounded up), for position relative to the Chili River (north/south) and urbanicity (urban/periurban). (B) District-level HIPSTR tree, with branches coloured by inferred district, and map of inferred movements between districts, with the Chili River shown in blue. Circle size shows time spent in each district (%), arrow width shows mean Markov jumps, and line type shows Bayes factor support for the route. Solid lines indicate decisive support and dashed lines very strong support for the route. Only transitions with Bayes factor > 50 are shown. The Chili River is shown in blue.

This structure held across broader divisions. Lineages were located mainly north of the river (69% of total evolutionary time vs 31% south), with bidirectional movement but a bias towards north-to-south transmission (13 vs 4 MJ). Urban areas were the primary foci (57% of evolutionary time), with substantially higher urban-to-periurban than periurban-to-urban transmission (mean 45 vs 12 MJ). Both directional biases were retained after downsampling (Table S3), although urban-to-periurban MJs fell from 45 to 25. This likely reflects densely sampled districts contributing disproportionately to inferred transitions, as in the district-level downsampling analysis.

No landscape variable was associated with lineage velocity (Q), but several were associated with viral persistence (E) and movement (R). Lineages were strongly associated with inhabited areas and higher human population density (decisive BF support). Relative population change, but not absolute population change or density, strongly facilitated dispersal, suggesting preferential movement towards expanding periurban areas.

Socioeconomic structure had a pronounced effect. Affluent areas consistently acted as barriers to both lineage occurrence and dispersal, whereas deprived areas were associated with greater persistence. Reversing the socioeconomic resistance surface did not produce symmetric effects, indicating that the spatial clustering of socioeconomic classes constrains diffusion in a non-reciprocal way. A continuous socioeconomic gradient gave weaker, less consistent effects.

Infrastructure also influenced movement. Major roads strongly facilitated dispersal, whereas travel time acted as a barrier. Water channels were associated with greater lineage occurrence but weaker effects on directed movement, and rivers were moderate partial barriers that constrained but did not prevent dispersal.

## Discussion

By integrating whole-genome sequencing with spatial and environmental data, we reconstructed a decade-long canine rabies epidemic in Arequipa, the first documented urban reintroduction in Latin America^16^. Rabies persisted primarily through sustained local transmission from a single lineage, concentrated in a few interconnected urban hubs and shaped by socioeconomic and landscape factors, rather than through repeated introductions. Much of this transmission was hidden from routine surveillance.

Introductions were nonetheless important. Despite regional connectivity within the Cosmopolitan AM5 subclade^10,28^, a single introduction seeded and subsequently dominated transmission in Arequipa city. This is consistent with observations from other endemic settings, including Harare, Zimbabwe^29^ and Gauteng Province, South Africa^30^ but notably different from other settings with multiple co-circulating lineages^7^. These differences likely reflect context-specific epidemiological and ecological factors, including vaccination coverage and its spatial distribution, regional endemicity and connectivity, and the structure and dynamics of local dog populations. Such factors may influence both the frequency with which new lineages are introduced and their potential to become established, shaping the extent to which local transmission is sustained by single or multiple co-circulating lineages^6,7,13^. Most other introductions, and likely others we did not detect, were epidemiological dead ends, but the 2021 El Pedregal outbreak shows that they can seed local transmission^10^. Bolivia has been repeatedly implicated as a source of canine rabies in Latin America^3,31^, where comparatively fragile control programmes have contributed to persistence^3,31^. These findings are consistent with metapopulation dynamics in which local transmission is sustained by occasional long-distance dispersal^32^.

The scale of the hidden transmission has implications for verifying elimination. We infer several years of cryptic transmission before first detection, and our outbreak size estimates indicate substantial under-detection. Where surveillance captures only a small fraction of cases, falling case counts are an unreliable indicator of elimination. Consistent with this, the estimated effective population size expanded rapidly during the early outbreak and then plateaued rather than declining. Although uncertainty increases towards the present, with wider credible intervals in recent estimates, the lack of a marked reduction in N1.τ despite declining reported cases (two cases in 2025 and none to date in 2026) highlights the difficulty of interpreting apparent reductions in incidence in settings with limited surveillance sensitivity and spatially heterogeneous transmission. Historical eradication efforts, most notably smallpox, illustrate that verification of elimination requires intensive, multi-layered surveillance rather than reliance on case reporting alone^33^. Here, genomic data added a complementary layer that provided key insights – dating the lineage’s emergence years before the first detected case and allowed us to separate sustained local transmission from introductions, neither of which case reports could show.

The hubs we identified, Cerro Colorado and neighbouring Yura, were previously noted as rabies hotspots characterised by socioeconomic disadvantage, reduced access to health services, and rapid urban expansion from migrant labour^19,34–37^. Our landscape analyses suggest that population and socioeconomic characteristics shape where transmission is maintained, whereas the urban landscape shapes how virus moves between areas. Affluent neighbourhoods may impede spread through greater confinement and vaccination of owned dogs, whereas deprived communities may sustain transmission through larger populations of free-roaming, unvaccinated dogs, and rapidly expanding peri-urban areas may seed new transmission chains^34^. The effect of roads is consistent with human-mediated dog movement along established transport corridors, as reported in other settings^7,8^. Here, we show that these landscape and socioeconomic effects also operate at fine intra-urban resolution. This reinforces calls for systems-oriented approaches that combine urban planning, dog population management, vaccination, and public health interventions^38,39^.

Latin America is the world’s most urbanised region, and cities such as Arequipa have undergone rapid expansion over recent decades. This is accompanied by increased migration, informality, and inequality, which has been associated with increased disease prevalence^34,40,41^. At a coarse scale, viral movement in Arequipa was predominantly from urban to peri-urban areas, but this should be interpreted cautiously. Binary classifications fail to capture urban heterogeneity^39^ and previous studies in Arequipa suggest that peri-urban settlements, including free-roaming and so-called feral or semi-feral “cave” dogs, may act as reservoirs^42^. Although truly feral dogs typically represent only a small proportion of free-roaming dog populations in most settings^43,44^, the cave-dog ecology described in Arequipa may be a localised exception^34,42^ and these dogs present a particularly challenging demographic to reach through conventional rabies control interventions. Lower vaccination coverage, greater roaming ranges and higher densities may allow these dogs to contribute disproportionately to transmission^34,42^. Limited sampling of them may bias our directionality estimates. In contrast, our finer-scale analyses suggest that transmission is structured more by intra-urban connectivity than by broad urbanisation gradients. The microred structure (sub-district operational units reflecting the organisation of community healthcare) may capture these social and spatial pathways better than binary classifications or administrative boundaries, and should inform future analyses and control, although we did not test this directly.

This study has limitations. Although this is the largest genomic dataset for canine rabies in the region, sampling is incomplete and uneven, with only 32 publicly available genomes from outside Peru. While this level of sampling is comparable to that of other WGS phylogeographic studies of RABV^6,7,13^, it reflects the broader scarcity of genomic data for this pathogen, and limits the resolution with which transmission pathways, directionality, and source-sink dynamics can be inferred. Although continuous phylogeographic approaches are generally less sensitive to sampling bias than discrete models^45–47^ and we took steps to test the robustness of our results, residual data bias (e.g. unsampled cave dogs) cannot be excluded^45^, and occasional long branches in our phylogeny also suggest unsampled transmission. Key results were nevertheless robust to downsampling. Sparse sampling also limits source attribution: sample 219_2024 clustered with sequences from Argentina, Bolivia, and Brazil, and epidemiological evidence was needed to infer a Bolivian origin, while 75_2021 clustered with historical Puno sequences, so a recent introduction cannot be distinguished from persistence in an unsampled chain. Since Puno Region has reported no canine or human rabies since 2023, this uncertainty raises the possibility of undetected circulation and highlights a priority for surveillance. Genomic epidemiology can resolve transboundary transmission, but only with more geographically representative sequencing. PAHO’s Cooperation Among Countries for Health Development initiative, strengthening joint Peru-Bolivia surveillance, offers an opportunity to build this capacity.

In summary, our findings have three implications for rabies elimination in urban endemic settings. First, case counts alone cannot establish elimination, and genomic data are needed to distinguish introductions from undetected persistence. Second, transmission is highly spatially heterogeneous, so intervention should be intensified in urban hubs and the corridors linking them. Third, recurrent introductions mean control must extend beyond the city, with coordinated regional surveillance. Although no cases have been detected so far in 2026 in Arequipa, the repeated introductions we identified, and the likelihood of undetected circulation, mean the risk of reintroduction remains high. Vaccination must therefore be maintained to prevent a new introduction from re-establishing transmission. Rabies elimination will require integrated control across scales, and the priority now is to use this evidence to target limited resources where they are likely to have the greatest effect.

## Acknowledgments

We thank the field and outbreak control teams from the One Health Unit at Universidad Peruana Cayetano Heredia and the Arequipa Health Network (Red de Salud Arequipa-Caylloma). We thank Dr. Ballón and his team at Universidad Nacional San Agustín (UNSA). We also thank the laboratory team at the Arequipa Regional Reference Laboratory (GERESA-LRR). Special thanks to Drs. Sergio Recuenco, Ana María Navarro, and Ricardo López-Ingunza for sharing valuable insights.

## Author contributions

Conceptualization: KB, ADB, RCN Methodology: KB, RD

Software: SK, KB Validation: ADB Formal analysis: KB, RD

Investigation: ADB, EWD, EZ Resources: KB, EZ, YM, RCN

Data Curation: KB, ADB, EWD, EZ Writing - Original Draft: KB, ADB, RCN Writing - Review & Editing: Visualization: KB

Supervision: KB, RCN

Project administration: KB, RCN Funding acquisition: KB, RCN

## Supplementary Information

Supplementary text, tables and figures are available in the Supplementary Information file.

## Data and code availability

Consensus genome sequences generated in this study will be deposited in GenBank prior to publication, with accession numbers to be provided upon submission. All code, analysis workflows, and supporting data required to reproduce the analyses are available at: https://doi.org/10.5281/zenodo.23808245^48^

