## Supplementary Material for "One Health genomic epidemiology reveals cryptic transmission and surveillance gaps in a decade-long urban rabies epidemic in Arequipa, Peru"

### Supplementary Information

#### Supplementary Text:

**Study grid**

A uniform spatial grid was constructed to define the study landscape for resistance surface analyses. All spatial data were first projected to a common coordinate reference system (UTM Zone 19S; EPSG:32719) to ensure that distances and areas were expressed in metres. Arequipa city case locations were used to define the spatial extent of the study area. To minimise edge effects in subsequent landscape analyses, each extent was expanded by a 1km buffer beyond the outermost case locations, resulting in a raster grid covering ~650km$^2$.

A regular lattice of square grid cells was then generated over each buffered extent using a fixed spatial resolution of 200 m x 200 m. This resolution was selected to balance computational tractability with the spatial scale of urban heterogeneity relevant to dog movement and transmission processes. Each grid cell was assigned a constant baseline value of 1, representing homogeneous space prior to incorporation of environmental resistance or conductance layers.

The resulting raster grid was implemented using the terra package in R, ensuring consistent alignment of all subsequent environmental layers. The grid was exported for downstream phylogeographic analyses in ASCII and GeoTIFF formats compatible with SERAPHIM.

**Landscape variables (resistance surface construction)**

Landscape predictors were derived from spatial covariates representing infrastructure, topography, socioeconomic structure, and population dynamics, and converted into continuous resistance surfaces aligned to the study grid. Sources for these data are in Table S1. All layers were projected to UTM Zone 19S (EPSG:32719) and rasterised to a 200 m x 200 m grid matching the base analysis surface above.

***Infrastructure surfaces***

Vector-based infrastructure data (roads, rivers, and water channels) were converted to spatial line features and rasterised onto the study grid. Only major roads and rivers (i.e. Chili River) were considered. For each feature class, two alternative representations were generated to capture different hypotheses about movement behaviour: (i) binary surfaces, in which grid cells intersecting a feature were assigned a value of 1 and all other cells 0, and (ii) distance-decay surfaces, where Euclidean distance to the nearest feature was calculated and converted into a declining influence function of the form $1 / (1 + d/\sigma)$, with $\sigma = 100$ m for urban-scale diffusion. All surfaces were normalised to a 0–1 range prior to scaling.

Roads and water channels were treated as potential facilitators of movement under distance-decay formulations and as discrete conduits under binary formulations. Rivers were additionally modelled as both potential barriers and conduits depending on parameterisation, reflecting uncertainty in their role as movement constraints in the urban environment.

***Habitation and Socioeconomic structure***

Habitation was represented as a binary spatial layer indicating the presence or absence of residential areas (houses). This surface was derived by rasterising habitation footprints onto the study grid, with occupied cells assigned a k-scaled resistance value (1 + k, while all other cells were assigned a baseline value of 1. This layer was used as a proxy for domestic dog presence and human–dog contact potential.

Socioeconomic status (SES) was derived from administrative micro-area classifications (A–F) from Xie et al. (2025) and converted into ordinal surfaces. Two formulations were used: (i) a continuous gradient reflecting decreasing socioeconomic status, and (ii) a binary classification contrasting affluent (A–C) and deprived (D–F) areas. Higher deprivation was treated as higher resistance under the assumption of increased vulnerability and reduced access to vaccination and dog confinement practices. Raster cells outside SES polygons were assigned a penalty value to avoid artefacts from missing data.

***Land cover (cropland)***

Cropland was derived from the Sentinel-2 10 m land use/land cover classification product (Esri and Impact Observatory), produced from Sentinel-2 imagery. The raster was reclassified to extract cropland areas and aligned to the study grid. Cropland was modelled as a binary resistance surface representing presence versus absence of agricultural land. Grid cells classified as cropland were assigned a k-scaled resistance value (1 + k), while all other cells were assigned a baseline value of 1. This formulation reflects the hypothesis that cropland may alter movement behaviour and human-animal interaction intensity, with sensitivity to effect strength evaluated across k.

***Population dynamics***

Built environment and population structure were represented using gridded population surfaces for 2015 and 2025, harmonised to the same spatial resolution and extent. Population density was treated as a proxy for host availability and modelled as increasing resistance under the assumption that higher host density facilitates transmission.

Temporal change in population density was derived using log-transformed differences between 2015 and 2025 surfaces to capture urban expansion dynamics. Alternative formulations included absolute change, relative (log-ratio) change, and z-score standardised change to assess robustness to scaling assumptions.

#### *Accessibility surface*

A gridded accessibility surface (50 m resolution) was incorporated as a proxy for transport connectivity and urban permeability. The layer was aligned to the study grid through cropping, reprojection, and resampling to match the 200 m analysis resolution.

#### *Standardisation and export*

All environmental layers were standardised to a common 0–1 scale and converted into resistance surfaces using a consistent SERAPHIM-compatible transformation:

$$vt = 1 + k \times v_0$$

where $v_0$ is the normalised environmental surface and $k \in \{10, 100, 1000\}$ controls the magnitude of landscape effect. This transformation was applied uniformly across all predictor layers to ensure comparability and to enable sensitivity analysis of effect strength across alternative parameterisations.

All final raster layers were exported in ASCII format for downstream phylogeographic analyses in SERAPHIM.

### Supplementary Tables:

**Table S1.** Data sources for landscape predictor variables.

| Predictor | Dataset / product | Provider | Reference | Access URL |
|---|---|---|---|---|
| Accessibility | Global accessibility (travel time to nearest city ≥50,000 population) | European Commission Joint Research Centre (JRC), Global Environment Monitoring Unit | [1] | https://forobs.jrc.ec.europa.eu/gam/download |
| Population density | Gridded population | WorldPop, University of Southampton | [2] | https://data.humdata.org/dataset/worldpop-population-counts-2015-2030-per |

| | | | | |
|---|---|---|---|---|
| | counts (2015–2030, R2025A v1) | | | |
| Socioeconomic status | High-resolution socioeconomic micro-area classifications (A–F) derived from Peru National Institute of Statistics and Informatics (INEI) census data. | University of Pennsylvania/ Universidad Peruana Cayetano Heredia Zoonotic Disease Research Centre (ZDRC) | [3,4] | https://doi.org/10.5281/zenodo.21398304<br>Derived from:<br>https://www.inei.gob.pe/media/MenuRecursivo/publicaciones_digitales/Est/Lib1747/libro.pdf |
| Habitation | Residential footprint / habitation layer (derived) | ZDRC | [3] | https://github.com/RabiesLabPeru/arequipa_spatialdata |
| Water channels | Urban water channels | ZDRC | [3,5] | https://github.com/RabiesLabPeru/arequipa_spatialdata |
| Cropland (land cover) | Sentinel-2 10 m Land Use/Land Cover classification | Esri & Impact Observatory (derived from ESA Copernicus Sentinel-2 imagery) | [6] | https://livingatlas.arcgis.com/landcoverexplorer/ |

| | | | | |
|---|---|---|---|---|
| | (cropland class extracted) | | | |
| Roads | Road network | ZDRC | 3 | https://github.com/RabiesLabPeru/arequipa_spatialdata |
| Rivers | River network | ZDRC | 3 | https://github.com/RabiesLabPeru/arequipa_spatialdata |

**Table S2. Sequencing and genome coverage metrics for the newly sequenced samples (n=133) included in this study.**

| Accession | Sample ID | Mapped reads | Mean depth (sd) | % Genome coverage |
|---|---|---|---|---|
| PZ671774 | 10_2020 | 102208 | 3691 (2405) | 99.19 |
| PZ671751 | 10_2021 | 170765 | 6327 (3939) | 99.19 |
| PZ671755 | 100_2023 | 73310 | 2655 (2102) | 99.19 |
| PZ671708 | 101_2017 | 21278 | 1114 (601) | 99.19 |
| PZ671718 | 103_2022 | 22171 | 1040 (684) | 99.19 |
| PZ671770 | 112_2017 | 100067 | 4035 (2851) | 99.19 |
| PZ671736 | 114_2015 | 7494 | 457 (207) | 99.19 |
| PZ671790 | 12_2017 | 9712 | 414 (303) | 99.19 |
| PZ671696 | 130_2016 | 122127 | 4873 (3970) | 99.19 |
| PZ671725 | 131_2019 | 130354 | 5774 (3394) | 99.19 |
| PZ671727 | 145_2017 | 92869 | 3679 (2758) | 99.19 |
| PZ671702 | 146_2021 | 181507 | 6440 (4149) | 99.19 |
| PZ671782 | 149_2020 | 65078 | 3005 (1853) | 99.19 |
| PZ671698 | 15_2021 | 132244 | 5259 (3117) | 99.19 |
| PZ671717 | 151_2024 | 61144 | 2702 (1601) | 99.19 |
| PZ671772 | 152_2023 | 45626 | 2213 (1123) | 99.19 |
| PZ671798 | 159_2017 | 14897 | 858 (476) | 99.19 |
| PZ671786 | 16_2021 | 22435 | 976 (614) | 99.19 |
| PZ671808 | 162_2018 | 30203 | 1135 (664) | 99.19 |
| PZ671777 | 162_2024 | 28520 | 1190 (615) | 99.19 |
| PZ671783 | 166_2016 | 163646 | 6531 (4150) | 99.19 |
| PZ671803 | 175_2023 | 101661 | 3544 (2558) | 99.19 |
| PZ671740 | 183_2019 | 64086 | 2120 (1716) | 99.19 |
| PZ671726 | 183_2023 | 99340 | 3813 (2349) | 99.19 |
| PZ671809 | 186_2018 | 114508 | 4359 (3050) | 99.19 |
| PZ671759 | 2_2017 | 15248 | 615 (381) | 99.19 |
| PZ671749 | 201_2017 | 185489 | 8390 (5685) | 99.19 |

| | | | | |
|---|---|---|---|---|
| PZ671693 | 209_2018 | 62872 | 2061 (1758) | 99.19 |
| PZ671752 | 218_2023 | 138447 | 5541 (3518) | 99.19 |
| PZ671760 | 219_2024 | 39237 | 2021 (1208) | 99.19 |
| PZ671741 | 22_2016 | 30597 | 1097 (892) | 99.19 |
| PZ671687 | 22_2018 | 35246 | 1349 (805) | 99.19 |
| PZ671707 | 225_2017 | 109652 | 4996 (2671) | 99.19 |
| PZ671681 | 225_2019 | 73067 | 2860 (2096) | 99.19 |
| PZ671729 | 225_2023 | 40544 | 1913 (1210) | 99.19 |
| PZ671703 | 227_2019 | 3927 | 244 (97) | 99.19 |
| PZ671712 | 234_2016 | 46268 | 2306 (1207) | 99.19 |
| PZ671720 | 236_2019 | 34318 | 1438 (1052) | 99.19 |
| PZ671695 | 24_2019 | 50659 | 2468 (1366) | 99.19 |
| PZ671700 | 240_2019 | 90112 | 3024 (2748) | 99.19 |
| PZ671714 | 240_2023 | 83845 | 3499 (1800) | 99.19 |
| PZ671748 | 242_2019 | 93109 | 3355 (2387) | 99.19 |
| PZ671709 | 244_2019 | 106659 | 4054 (3344) | 99.19 |
| PZ671779 | 246_2022 | 69980 | 3583 (1844) | 99.19 |
| PZ671685 | 247_2018 | 86494 | 3402 (2414) | 99.19 |
| PZ671806 | 255_2019 | 70841 | 3035 (2502) | 99.19 |
| PZ671737 | 259_2017 | 88256 | 4614 (2563) | 99.19 |
| PZ671722 | 266_2022 | 62829 | 2593 (1842) | 99.19 |
| PZ671738 | 270_2023 | 25459 | 1033 (566) | 99.19 |
| PZ671795 | 272_2019 | 31950 | 1328 (649) | 99.19 |
| PZ671704 | 273_2023 | 93859 | 3652 (2192) | 99.19 |
| PZ671804 | 275_2017 | 113006 | 5394 (3226) | 99.19 |
| PZ671694 | 280_2022 | 22516 | 1290 (678) | 99.19 |
| PZ671780 | 289_2019 | 17211 | 699 (499) | 99.19 |
| PZ671799 | 29_2021 | 148676 | 5826 (3667) | 99.19 |
| PZ671678 | 297_2023 | 57856 | 2185 (1704) | 99.19 |
| PZ671789 | 298_2017 | 33064 | 1205 (754) | 99.19 |
| PZ671778 | 302_2016 | 35354 | 1854 (822) | 99.19 |
| PZ671762 | 302_2017 | 260343 | 11226 (7998) | 99.19 |
| PZ671733 | 312_2019 | 41694 | 1596 (1054) | 99.19 |
| PZ671756 | 315_2017 | 15145 | 631 (384) | 99.19 |
| PZ671768 | 32_2016 | 38464 | 1663 (834) | 99.19 |
| PZ671692 | 320_2019 | 182795 | 6169 (4010) | 99.19 |
| PZ671747 | 333_2018 | 74700 | 3178 (2525) | 99.19 |
| PZ671734 | 348_2023 | 75768 | 2966 (1603) | 99.19 |
| PZ671689 | 35_2018 | 19014 | 1346 (540) | 99.19 |
| PZ671711 | 357_2016 | 179013 | 8311 (5210) | 99.19 |

| | | | | |
|---|---|---|---|---|
| PZ671802 | 358_2023 | 147444 | 5869 (3583) | 99.19 |
| PZ671784 | 367_2015 | 35263 | 1853 (984) | 99.19 |
| PZ671792 | 374_2018 | 48191 | 2468 (1040) | 99.19 |
| PZ671683 | 375_2018 | 18425 | 1049 (448) | 99.19 |
| PZ671723 | 386_2016 | 106814 | 4356 (2586) | 99.19 |
| PZ671800 | 4_2023 | 9229 | 463 (271) | 99.19 |
| PZ671754 | 402_2017 | 13176 | 894 (437) | 99.19 |
| PZ671701 | 41_2024 | 28426 | 1414 (629) | 99.19 |
| PZ671791 | 435_2018 | 90938 | 3808 (2861) | 99.19 |
| PZ671793 | 44_2016 | 80975 | 3571 (2070) | 99.19 |
| PZ671761 | 442_2016 | 115899 | 4444 (2941) | 99.19 |
| PZ671677 | 447_2018 | 110154 | 4342 (3062) | 99.19 |
| PZ671686 | 46_2017 | 31755 | 1715 (739) | 99.19 |
| PZ671699 | 46_2019 | 61402 | 2672 (1661) | 99.19 |
| PZ671735 | 462_2016 | 28902 | 1262 (815) | 99.19 |
| PZ671730 | 468_2016 | 30965 | 1244 (696) | 99.19 |
| PZ671743 | 48_2019 | 141184 | 7348 (4425) | 99.19 |
| PZ671697 | 480_2016 | 154408 | 6213 (4231) | 99.19 |
| PZ671732 | 481_2016 | 37121 | 1903 (1051) | 99.19 |
| PZ671766 | 485_2016 | 46423 | 2435 (1218) | 99.19 |
| PZ671684 | 495_2016 | 144640 | 5757 (4260) | 99.19 |
| PZ671788 | 51_2018 | 57187 | 2166 (1566) | 99.19 |
| PZ671807 | 54_2024 | 21647 | 1032 (443) | 99.19 |
| PZ671706 | 56_2023 | 65574 | 2597 (1636) | 99.19 |
| PZ671682 | 562_2016 | 120625 | 4693 (3115) | 99.19 |
| PZ671742 | 563_2016 | 91188 | 3682 (2979) | 99.19 |
| PZ671801 | 57_2017 | 208113 | 8861 (5778) | 99.19 |
| PZ671797 | 571_2016 | 30293 | 1445 (1093) | 99.19 |
| PZ671719 | 576_2016 | 118001 | 5042 (3163) | 99.19 |
| PZ671773 | 609_2015 | 32485 | 1345 (910) | 99.19 |
| PZ671680 | 653_2015 | 32909 | 1370 (659) | 99.19 |
| PZ671776 | 704_2015 | 39784 | 1662 (1065) | 99.19 |
| PZ671690 | 711_2015 | 104192 | 3578 (2573) | 99.19 |
| PZ671758 | 73_2015 | 45067 | 2319 (1160) | 99.19 |
| PZ671765 | 89_2018 | 109076 | 3686 (2956) | 99.19 |
| PZ671775 | 9_2019 | 254819 | 10567 (7991) | 99.19 |
| PZ671739 | 91_2017 | 154210 | 6917 (4699) | 99.19 |
| PZ671753 | 92_2018 | 27450 | 1380 (598) | 99.19 |
| PZ671710 | 97_2017 | 49354 | 2067 (1099) | 99.19 |
| PZ671744 | 291_2016 | 18538 | 810 (670) | 99.01 |

| | | | | |
|---|---|---|---|---|
| PZ671721 | 534_2016 | 61995 | 2093 (1629) | 98.94 |
| PZ671769 | 213_2021 | 19107 | 666 (459) | 97.22 |
| PZ671767 | 272_2017 | 23174 | 883 (800) | 97.22 |
| PZ671750 | 42_2020 | 8194 | 304 (208) | 97.22 |
| PZ671785 | 47_2019 | 29480 | 1049 (731) | 97.22 |
| PZ671794 | 69_2016 | 20340 | 744 (475) | 97.22 |
| PZ671757 | 140_2018 | 40863 | 1485 (1175) | 97.14 |
| PZ671679 | 232_2023 | 39621 | 1708 (928) | 97.14 |
| PZ671728 | 286_2022 | 32778 | 1338 (1134) | 97.14 |
| PZ671713 | 88_2019 | 40894 | 1495 (1260) | 97.14 |
| PZ671715 | 279_2017 | 43405 | 1540 (1158) | 96.94 |
| PZ671688 | 69_2022 | 47559 | 1829 (1421) | 96.94 |
| PZ671787 | 93_2018 | 6139 | 285 (177) | 95.20 |
| PZ671763 | 336_2019 | 68513 | 2301 (2015) | 95.03 |
| PZ671764 | 321_2023 | 74049 | 2445 (1803) | 94.89 |
| PZ671805 | 25_2020 | 15844 | 566 (527) | 94.54 |
| PZ671781 | 30_2023 | 18353 | 670 (645) | 94.54 |
| PZ671691 | 52_2018 | 4944 | 202 (146) | 94.54 |
| PZ671745 | 696_2015 | 26881 | 930 (784) | 94.54 |
| PZ671724 | 699_2015 | 28854 | 995 (796) | 94.54 |
| PZ671771 | 41_2020 | 9313 | 344 (287) | 92.36 |
| PZ671731 | 319_2017 | 12897 | 472 (411) | 92.35 |
| PZ671746 | 75_2021 | 11273 | 395 (352) | 88.33 |
| PZ671705 | 23_2020 | 6831 | 248 (220) | 86.55 |
| PZ671716 | 140_2020 | 12146 | 437 (559) | 84.92 |
| PZ671796 | 40_2025 | 24604 | 790 (749) | 84.19 |

**Table S3. Robustness of inferred discrete trait transmission pathways under downsampling.** Discrete phylogeographic transitions with strong support in the full dataset (BF > 10) were assessed for sensitivity to sampling intensity using Treemmer-based downsampling. District-level analyses were repeated using two independently downsampled datasets (100 sequences; minimum 10 sequences per district), while river and urbanicity analyses were evaluated using a single downsampled dataset. For each transition, Bayes factors (BF) and mean Markov jumps (MJ) are reported for the full and downsampled datasets. Transitions were classified as Robust when their directionality and statistical support were retained following downsampling, and as Sensitive when they were not consistently recovered.

| Trait | From | To | BF (Full) | BF (DS1) | BF (DS2) | MJ (Full) | MJ (DS1) | MJ (DS2) | Classification |
|---|---|---|---|---|---|---|---|---|---|
| District | ccolorado | yura | 119631.39 | 393.66 | 39.34 | 19.61 | 8.72 | 8.77 | Robust |

| | | | | | | | | | |
|---|---|---|---|---|---|---|---|---|---|
| District | mmelgar | ccolorado | 9957.1 | 77.07 | 138.16 | 3.98 | 3.95 | 3.89 | Robust |
| District | ccolorado | cayma | 1581.97 | 594.04 | 5.82 | 9.24 | 7.77 | 2.28 | Robust |
| District | mmelgar | miraflores | 1581.97 | 121.59 | 115.91 | 3.9 | 3.74 | 3.92 | Robust |
| District | asa | jlbyr | 1036.22 | 337.57 | 84.54 | 4.69 | 4.42 | 4.4 | Robust |
| District | ccolorado | asa | 142.7 | 22.97 | 37.11 | 7.03 | 5.12 | 6.52 | Robust |
| District | asa | hunter | 112.38 | 98.42 | 23.65 | 2.79 | 3.17 | 2.16 | Robust |
| District | yura | ccolorado | 57.29 | 2.27 | 11.63 | 3.66 | 0.24 | 2.53 | Robust |
| District | asa | mmelgar | 53.1 | 48.22 | 28.67 | 3.79 | 4.29 | 4.13 | Robust |
| District | mollebaya | elpedregal | 47.16 | 58.74 | 38.77 | 0.77 | 0.81 | 0.75 | Robust |
| District | yura | socabaya | 18.87 | 13.74 | 3.9 | 1.75 | 1.52 | 0.61 | Robust |
| District | yura | paucarpata | 17.77 | 31.57 | 10.67 | 0.98 | 1.1 | 0.66 | Robust |
| *District* | *mmelgar* | *asa* | *17.33* | *9.98* | *7.3* | *1.32* | *1.34* | *1.46* | *Sensitive* |
| District | mmelgar | paucarpata | 15.34 | 23.04 | 16.49 | 1.35 | 1.64 | 1.29 | Robust |
| District | ccolorado | uchumayo | 11.68 | 17.95 | 4.88 | 0.57 | 0.75 | 0.36 | Robust |
| *District* | *yura* | *hunter* | *11.56* | *6.47* | *3.62* | *0.99* | *0.48* | *0.42* | *Sensitive* |
| *District* | *asa* | *paucarpata* | *11.26* | *5.4* | *6.76* | *1.41* | *0.8* | *1.13* | *Sensitive* |
| River | north | south | 11050.89 | 11050.89 | - | 12.57 | 11.94 | - | Robust |
| River | south | external | 268.34 | 104.03 | - | 1.0 | 1.00 | - | Robust |
| River | south | north | 154.44 | 20.57 | - | 4.25 | 5.70 | - | Robust |
| Urbanicity | periurban | urban | 11050.89 | 2761.80 | - | 11.93 | 8.37 | - | Robust |
| Urbanicity | urban | periurban | 11050.89 | 11050.89 | - | 44.87 | 25.28 | - | Robust |
| Urbanicity | periurban | external | 41.61 | 48.56 | - | 0.97 | 0.98 | - | Robust |

**Table S4. Supported transitions across all discrete trait analyses.** Subset of transitions with Bayes factor support >10, indicating statistically supported viral movement across all discrete trait models.

| Trait | From | To | Posterior probability | Bayes factor | Support category | Mean Markov jumps |
|---|---|---|---|---|---|---|
| District | ccolorado | yura | 1 | 119631.39 | Decisive | 19.61 |
| District | mmelgar | ccolorado | 1 | 9957.1 | Decisive | 3.98 |
| District | ccolorado | cayma | 0.99 | 1581.97 | Decisive | 9.24 |
| District | mmelgar | miraflores | 0.99 | 1581.97 | Decisive | 3.9 |
| District | asa | jlbyr | 0.99 | 1036.22 | Decisive | 4.69 |
| District | ccolorado | asa | 0.91 | 142.7 | Decisive | 7.03 |
| District | asa | hunter | 0.89 | 112.38 | Decisive | 2.79 |
| District | yura | ccolorado | 0.81 | 57.29 | Very strong | 3.66 |
| District | asa | mmelgar | 0.8 | 53.1 | Very strong | 3.79 |
| District | mollebaya | elpedregal | 0.78 | 47.16 | Very strong | 0.77 |
| District | yura | socabaya | 0.59 | 18.87 | Strong | 1.75 |
| District | yura | paucarpata | 0.57 | 17.77 | Strong | 0.98 |
| District | mmelgar | asa | 0.57 | 17.33 | Strong | 1.32 |
| District | mmelgar | paucarpata | 0.54 | 15.34 | Strong | 1.35 |
| District | ccolorado | uchumayo | 0.47 | 11.68 | Strong | 0.57 |
| District | yura | hunter | 0.47 | 11.56 | Strong | 0.99 |
| District | asa | paucarpata | 0.46 | 11.26 | Strong | 1.41 |
| River | north | south | 1 | 11050.89 | Decisive | 6.28 |
| River | south | external | 1 | 268.34 | Decisive | 0.5 |
| River | south | north | 0.99 | 154.44 | Decisive | 2.13 |
| Urbanicity | periurban | urban | 1 | 11050.89 | Decisive | 11.93 |
| Urbanicity | urban | periurban | 1 | 11050.89 | Decisive | 44.87 |
| Urbanicity | periurban | external | 0.97 | 41.61 | Very strong | 0.97 |

**Table S5. Lineage persistence across discrete trait models.** Markov reward estimates and percentage lineage persistence for each state across river, urbanicity, and district models.

| Trait | State | Markov reward (Years) | Persistence (%) |
|---|---|---|---|
| District | ccolorado | 103.45 | 51.6 |
| District | mmelgar | 27.42 | 13.7 |

| | | | |
|---|---|---|---|
| District | yura | 26.51 | 13.2 |
| District | asa | 14.14 | 7.1 |
| District | cayma | 7.31 | 3.6 |
| District | socabaya | 3.95 | 2 |
| District | jlbyr | 3.59 | 1.8 |
| District | paucarpata | 3.15 | 1.6 |
| District | characato | 2.77 | 1.4 |
| District | miraflores | 2.73 | 1.4 |
| District | mollebaya | 1.81 | 0.9 |
| District | hunter | 1.67 | 0.8 |
| District | uchumayo | 1.17 | 0.6 |
| District | sachaca | 0.59 | 0.3 |
| District | elpedregal | 0.4 | 0.2 |
| River | north | 136.7 | 69.5 |
| River | south | 60.09 | 30.6 |
| River | external | 0.34 | 0.2 |
| Urbanicity | urban | 115.26 | 56.7 |
| Urbanicity | periurban | 88.01 | 43.3 |
| Urbanicity | external | 0.35 | 0.2 |

Supplementary Figures:

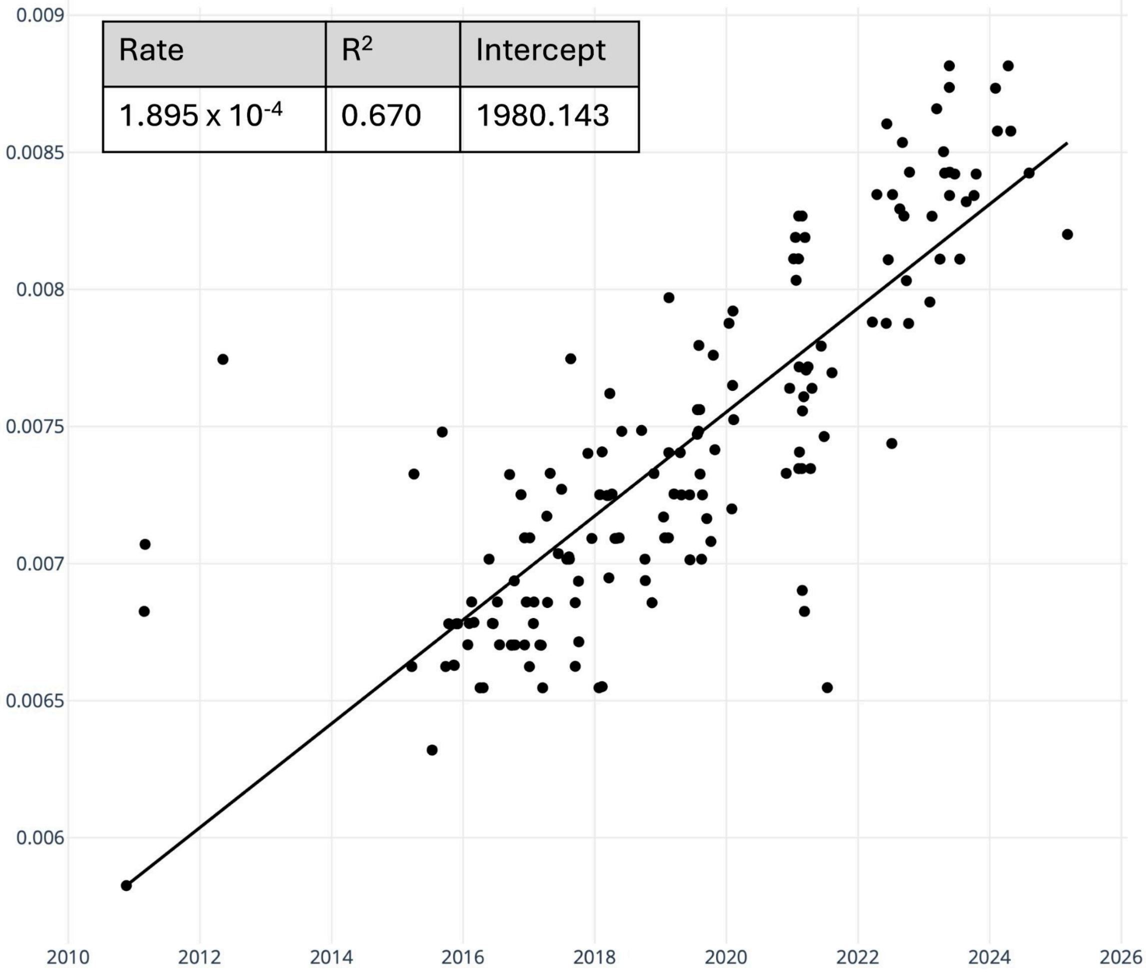


**Supplementary Fig. S1.** Root-to-tip genetic divergence against sampling date for Peruvian rabies virus genomes (n = 167), estimated in Clockor2. The regression line indicates temporal signal; inset shows $R^2$, substitution rate, and intercept.

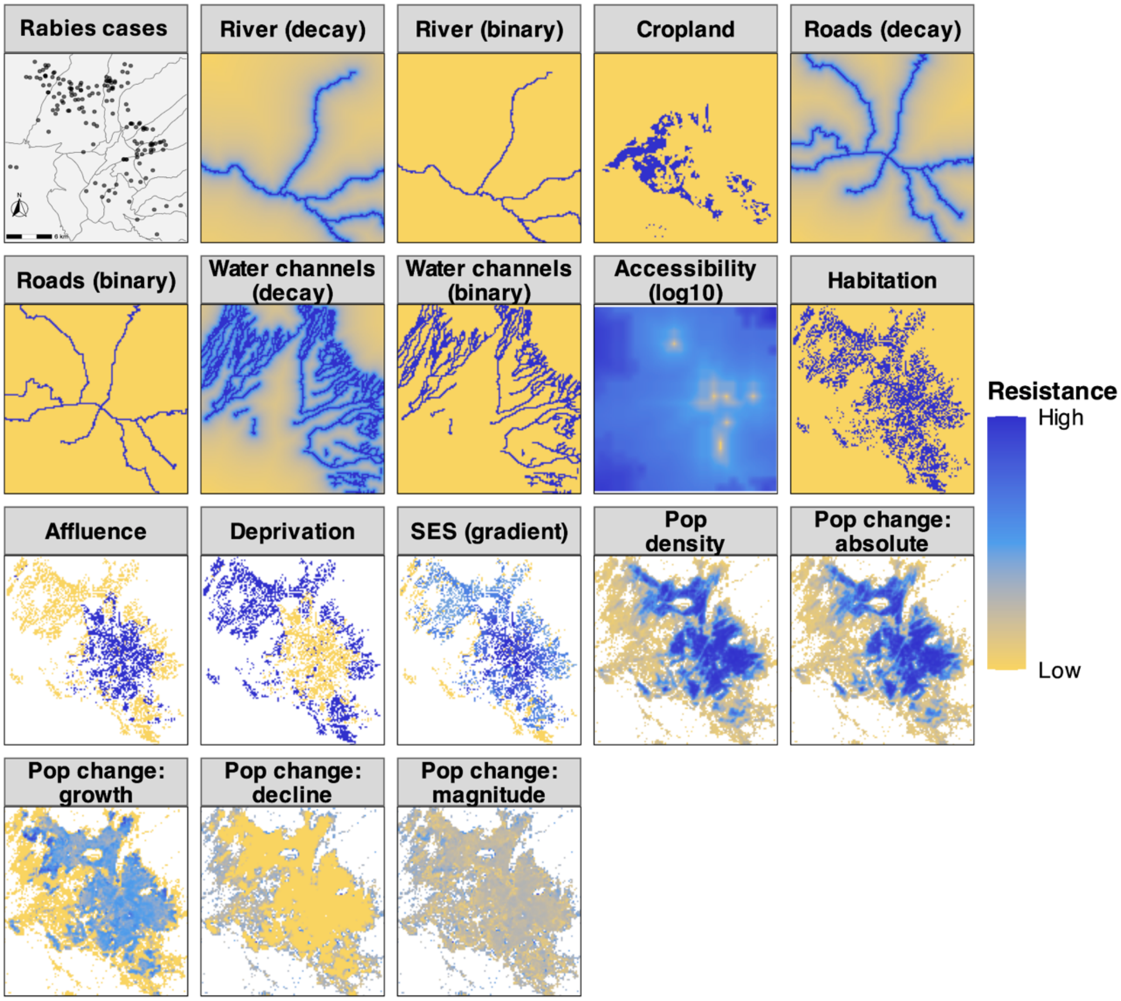


**Supplementary Fig. S2. Landscape variables used to model RABV spread in Arequipa, represented as resistance surfaces.** Variables include features representing movement barriers, land cover, accessibility, socio-economic context and population dynamics. The first panel shows the study area and distribution of sequenced rabies cases used in the analysis. Linear features (rivers, roads, and water channels) were represented as binary surfaces or distance-decay surfaces, where influence decreases with distance.

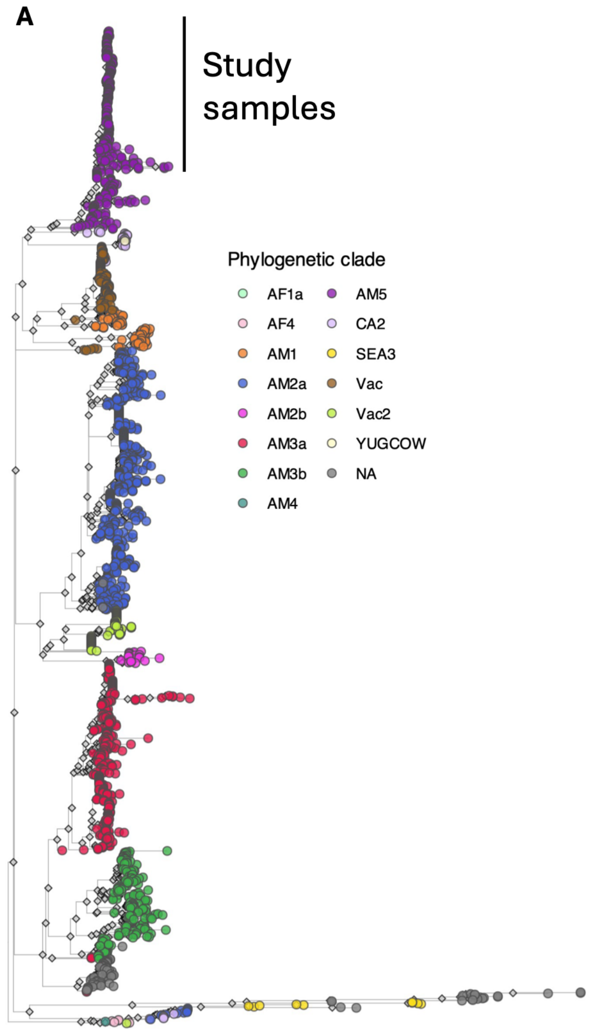


**Supplementary Fig. S3.** Maximum-likelihood phylogenetic trees of canine rabies virus sequences from Latin America and the Caribbean (LAC), including sequences of any available length (>200 bp). The dataset was assembled to provide regional-scale phylogenetic context and to identify dominant lineages circulating in Arequipa prior to lineage-specific subsetting (AM5-focused analyses presented in the main text). Trees are coloured by (A) phylogenetic minor clade and (B) country of origin. An outgroup representing the Cosmopolitan Africa 4 clade (GenBank: KF154998) was included for rooting but is not shown.
